# Assessing the effects of fertilisation on the yield, protein, and oil content of soybean seeds: a metadata analysis

**DOI:** 10.1101/2025.09.16.676653

**Authors:** Felipe Sousa Franco, Gustavo Paparotto Lopes, Nicolas Gustavo da Cruz da Silva, Julia Rossatto Brandão, Nícolas Leite Capucin, Felipe Rodrigues dos Santos, Gabriel Sgarbiero Montanha, Hudson Wallace Pereira de Carvalho

## Abstract

For a comprehensive study on balancing yield and grain composition, a meta-analysis of 48 peer-reviewed studies evaluated the effects of soil and foliar fertilization on soybean (*Glycine max* L.) yield, protein, and oil content. The metadata includes fertilizer type, application method, yield (kg ha^−1^), and concentration of protein (%) and oil (%). The data show that foliar and soil N fertilization increased grain yield by 13.7% and 14.2%, respectively (p < 0.05), compared to the control. However, it did not significantly affect the protein (38.4 ± 3.4%) or oil (20.2 ± 2.3%) content. High variability in experimental designs and the underrepresentation of foliar strategies make it difficult to draw broad conclusions. These findings showed the need for standardized field and controlled trials to balance increased yield with quality and to guide nutrient management.

## 1. SOYBEAN AS A STRATEGIC SOURCE OF PROTEIN AND OIL

Soybean (*Glycine max* (L.) Merrill) is a key source of food, feed, and raw chemicals for industry. Soybean seeds exhibit high protein and oil contents (*ca*. 30-42 and 18-22 wt.%, respectively), making them one of the most important oilseeds, as well as the biggest protein meal source for humans and animals worldwide (USDA, 2023).

Hence, the employment of agricultural strategies and management that seek to boost soybean seed quality, *i*.*e*., the content of relevant biomolecules, mainly protein and oil, receives increasing interest. Nevertheless, the biosynthesis of these molecules is tightly regulated through complex metabolic pathways, and most of the genetically based strategies aiming at increasing seed protein content resulted in low-yield accessions (USDA, 2020), which are economically unsustainable.

Notably, the massive efforts to increase yields over the last decades were followed by a consistent decline in seed protein concentration (Borja Reis et al., 2020). A comprehensive study on soybean varieties released in Brazil between 1965 and 2015 showed that the seed’s protein content exhibited a 1.1% decrease rate per decade (Umburanas et al., 2022), revealing that market-effective breeding programmes have not explored this trait.

Most of the soybean seed proteome is composed of seed storage proteins (SSPs), those to be used during seed germination and early seedling developmental stages (Montanha, G. S., 2025). In addition to genetic and environmental conditions, the availability of nutrients also plays a crucial role in the biosynthesis of storage proteins. Thus, to maximise the output-to-input ratio, in addition to yield gains, the effects of mineral nutrients on the content and quality of soybean seed proteins have drawn significant attention.

Among the mineral nutrients, sulfur (S) plays a crucial role in protein composition. Around 80% of S is found in protein in plants (Borja Reis et al., 2021). Additionally, soybeans naturally exhibit high levels of sulfur-containing amino acids (SAAs), *i*.*e*., as methionine and cysteine (Ma Y. et al., 2016).

A recent meta-analysis showed that S availability and assimilation by soybeans are intrinsically linked with the expression of SAAs in vegetative and reproductive organs (Hitsuda *et al*., 2015; Marschner, 2011). In this context, it was observed that applying S to soybean seeds at sowing led to a 0.3% increase in seed protein concentration, thereby potentially contributing to enhancing its nutritional quality through the increase of SAA relative abundance.

Moreover, soybean species are remarkably known for their ability to fix atmospheric nitrogen through a symbiotic association with *Bradyrhizobium* spp. bacteria, and hence, virtually no nitrogen (N) fertilisation is employed in most of the soybean-producing regions. Nevertheless, some pieces of evidence suggest that additional exogenous N could improve grain yield and seed quality (Wilson et al., 2014).

Altogether, these figures clearly indicate that a proper supply of mineral nutrients impacts both yield and biomolecules biosynthesis, there is a limited number of studies systematically assessing this topic, making such information virtually unavailable for farmers.

Therefore, the present study aims to determine the effects of application strategies of fertilisers on the quantitative and qualitative traits of soybean seeds. To address these questions, a metadata analysis was carried out from peer-reviewed publications found in the Web of Science Database.

## 2. EFFECTS OF MINERAL NUTRITION ON SOYBEAN YIELD, PROTEIN, AND OIL CONTENT

### 2.1 Experimental design and statistical analysis

The search in the Web of Science database was performed in December 2022 employing the following keywords and boolean operators: *“soybean” AND “fertilisation” AND “protein” OR “oil” AND “yield”* and returned 87 articles. After screening, only 48 articles reporting the specific quantitative data required for our analysis, *i.e*., exploring the effects of fertilisation *via* both soil and foliar application, on soybean seed yield and biomolecules (protein and oil) composition were selected for the metadata analysis. Each article was reviewed to retrieve information regarding i) the experimental strategy explored, i.e., whether the experiments were carried out under field, growth chamber, or greenhouse conditions; ii) the physicochemical properties of the soil used; iii) the nutrients applied, its dose and way of exposure, i.e., soil *or foliar application*; iv) the effects on agronomic parameters, i.e., yield, root and shoot length and dry mass, as well as seed weight, protein, and oil content. The compiled metadata information is freely available at the *figshare* repository (https://doi.org/10.6084/m9.figshare.30136411.v1).

### 2.2 Profile of fertiliser sources and application approaches

Figure 1 reveals that, from the 48 articles selected, which comprises 919 treatments, a major fraction of the studies (66.7% of the total) explored soil fertilisation strategies, whereas only 14.6% assessed the effects of foliar application (Figure 1a). Regarding the analysed attributes, it shows that most studies (56.3% of the total) reported effects on yield, protein, and oil content, whereas 23% evaluated either just yield or biomolecules, i.e., protein and oil contents (Fig. 1b).

**Figure 1.**
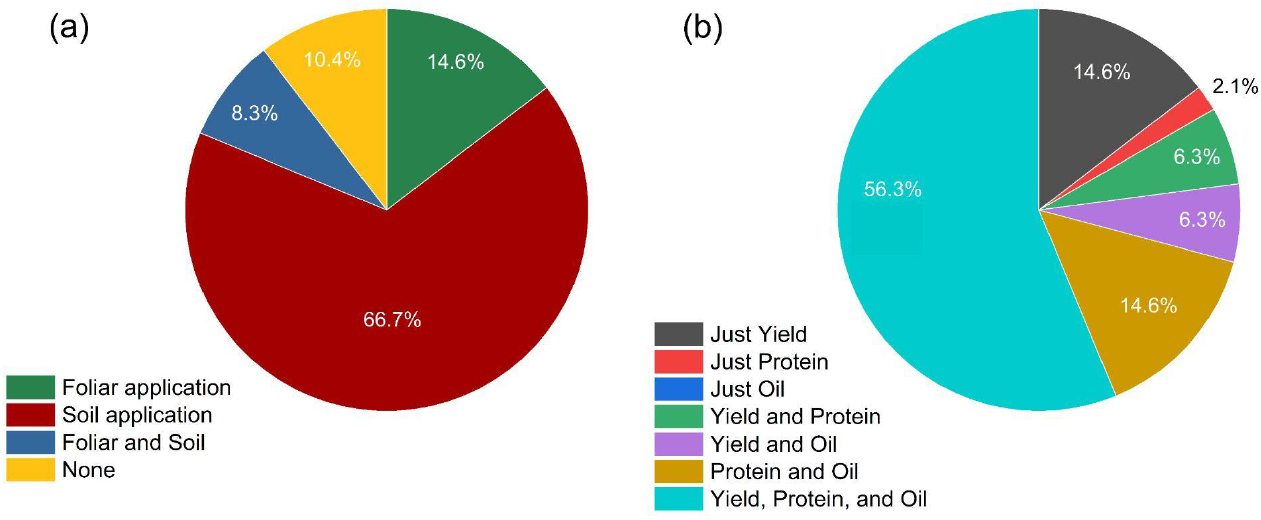
Quantitative overview on the fertilisation strategy, *i.e*., soil and/or foliar-based applications (a) and the agricultural method, i.e., seed yield and/or macromolecules (protein and oil) contents (b), reported in the 48 articles (n = 919 treatments) herein explored.

Moreover, the effects of soil and foliar-based fertilization on yield, protein, and oil contents are compared in Figure 2. Interestingly, significant increases in yields were obtained as a function of foliar fertilization compared to soil-based ones (3,853 and 3,313 kg ha^−1^, respectively), whereas no clear differences were observed for both oil and protein seed concentrations.

**Figure 2:**
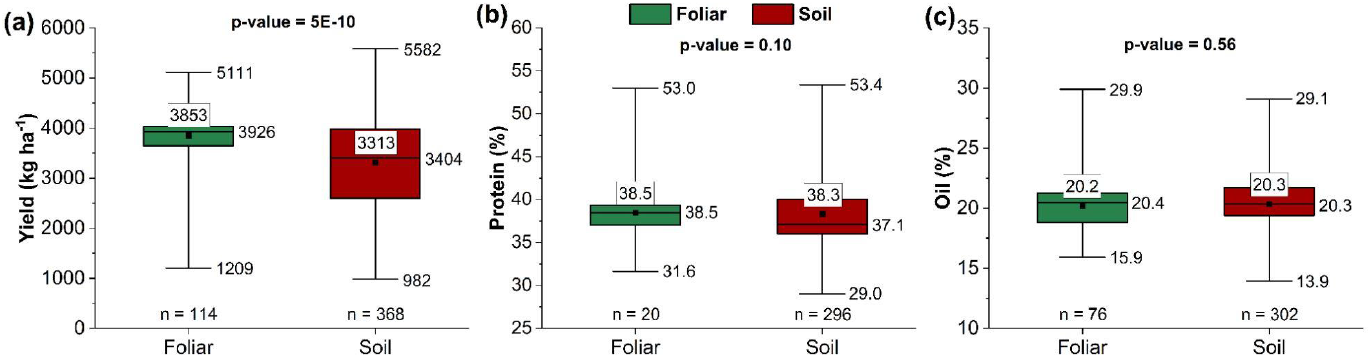
Effect of fertilizer application method, foliar or soil, on soybean yield (kg ha^−1^), protein content (%), and oil content (%). *p-value from Mann-Whitney test.

These numbers reinforce the importance of fertilization in increasing biomass production. Notably, it highlights the potential of foliar-deliver approaches over soil-based ones. Foliar fertilization stands out as a promising alternative for offering nutrients in high-demand periods, *e.g*. flowering or grain filling, as root absorption is not enough to achieve the high demand by modern crops (Meher, A. G. et al. 2024). Several studies have demonstrated the benefits of foliar micronutrient application in soybean, such as increased chlorophyll and carotenoid contents, which enhance the plant’s tolerance to environmental stresses (Falaknaz et al., 2022). This nutritional strategy reduces the plant’s vulnerability to climatic instabilities, *i.e*. drought conditions, by strengthening physiological responses, as increasing proline production (Zolfaghari Gheshlaghi, 2019).

Among the nutrients applied, micronutrients such as zinc (Zn), manganese (Mn), copper (Cu), and boron (B) are the most commonly used in foliar applications in soybean. This preference is mainly due to the challenges associated with their availability in the soil, which can be strongly influenced by redox conditions, soil texture (clay content), organic matter, pH, cation exchange capacity (CEC), and soil moisture. These factors can reduce the solubility and mobility of these elements in the rhizosphere, leading to partial unavailability to plants and potentially limiting productivity. Consequently, these micronutrients are among the most frequently studied in recent academic research on soybean nutrition (Ishfaq, M. et al., 2022).

Curiously, N, S, and K stand out as the most reported nutrients in the soil-based studies herein explored (Fig. 3a), whereas N, P, and K were mostly employed for soil-based ones (Fig. 3b). Despite the variability in the data, N was applied at higher rates compared to other nutrients in both soil and foliar strategies.

**Figure 3:**
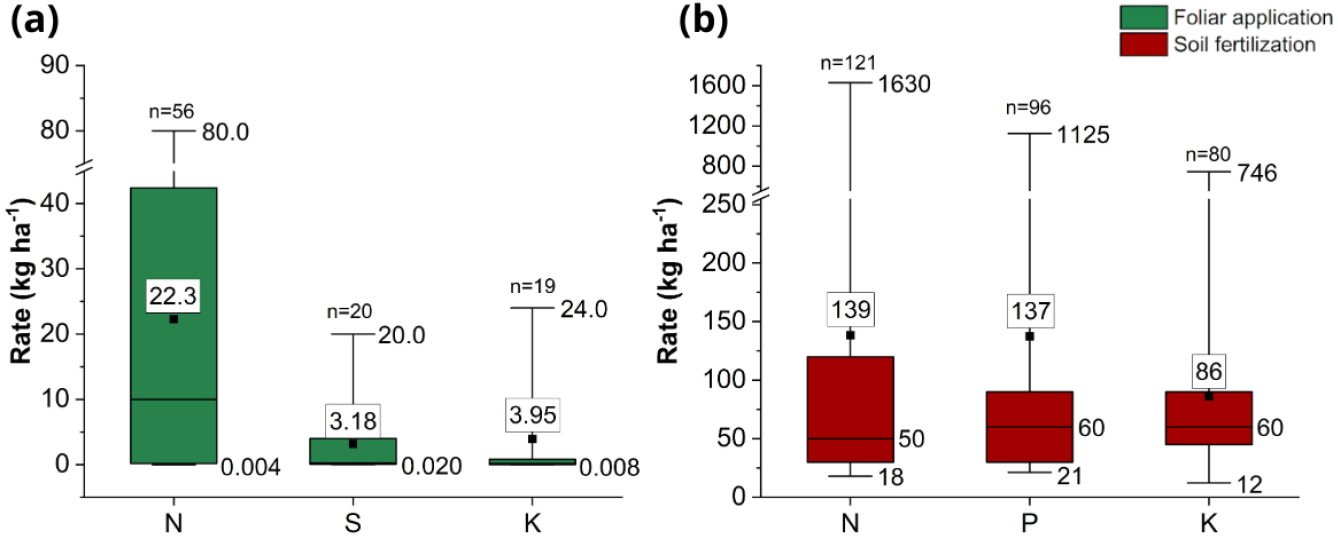
The different rates applied of each element (kg ha^−1^) by foliar and soil methods.

Furthermore, one should notice that N and K soil-applied (139 and 86 kg ha^−1^, respectively) doses were significantly higher than those applied via foliar (22.3 and 3.95 kg ha^−1^, respectively).

Considering the results shown in Figure 3, the foliar supply of macronutrients can offer several advantages in improving nutrient use efficiency during the plant’s high-demand developmental stages. Foliar absorption efficiency can reach up to 80%, compared to soil application, which often does not exceed 50% for certain nutrients due to environmental losses such as leaching and adsorption by clay particles (Fernández, V., & Brown, P. H., 2013). Moreover, soil applications are typically made only at the time of crop establishment, making late-season nutrient supply via soil unfeasible during critical periods of high physiological demand for grain production. Foliar application thus allows for better adjustment of the plant’s nutritional requirements, although it does not replace the base fertilization via soil. In this context, Dass *et al*. (2022) reported an increase in grain yield of approximately 18.6% to 20% following foliar application of a 2% urea solution. Similarly, Gowthami *et al*. (2018) demonstrated that combined foliar application of potassium and micronutrients can increase productivity by up to 28.5% compared to untreated controls in soybean fields.

### 2.3 Effects of fertilisation on grain quality

Figure 4 shows the reported data for yield, protein, and oil as a function of nutrient application (via soil or foliar). Once the Shapiro-Wilk test indicates that the data follows a non-normal distribution, the Mann-Whitney test was applied to compare the distributions between groups. The data revealed that nutrient application led to a statistically significant increase in the yield of 6.9%, whereas no changes were observed for both oil and protein content.

**Figure 4.**
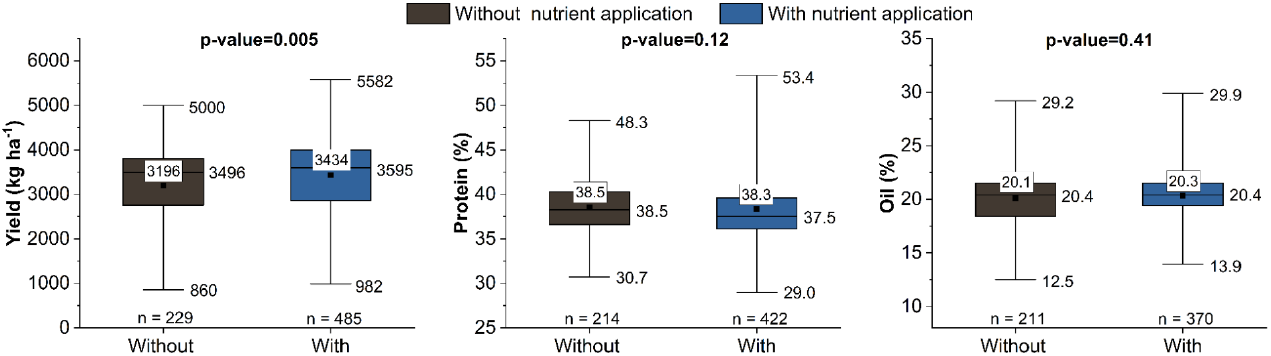
The correlation between yield (kg ha^−1^), protein (%) and oil (%) with the application or non-application of nutrients. *p-value from Mann-Whitney test.

This trend is highlighted by correlation analysis shown in the Supplementary Figure S4, which revealed only a slightly positive trend between grain yield, protein, and oil concentrations. On one hand, such values suggest that variations in yield are not strongly explained by changes in protein or oil content under the conditions evaluated in this metadata analysis. However, one should keep in mind that an increase in yields while maintaining its actual protein and oilseed content will lead to an increase in the amount of both biomolecules per unit of cultivated area. These results highlight the complexity of the physiological and environmental interactions that regulate soybean seed composition.

#### 2.2.3 Effects of nitrogen fertilization

Interestingly, more than half of the studies evaluated the application of N. Figure 5 shows the yield, protein, and oil content in studies that applied nitrogen compared to the control. A significant boost in yield (13.7%) was observed as a function of foliar N fertilisation. Additionally, differences were noted in grain yield, with an increase of 14.2% in N application treatments and a reduction of 2.9% in oil content when N was applied via soil compared to the control.

**Figure 5:**
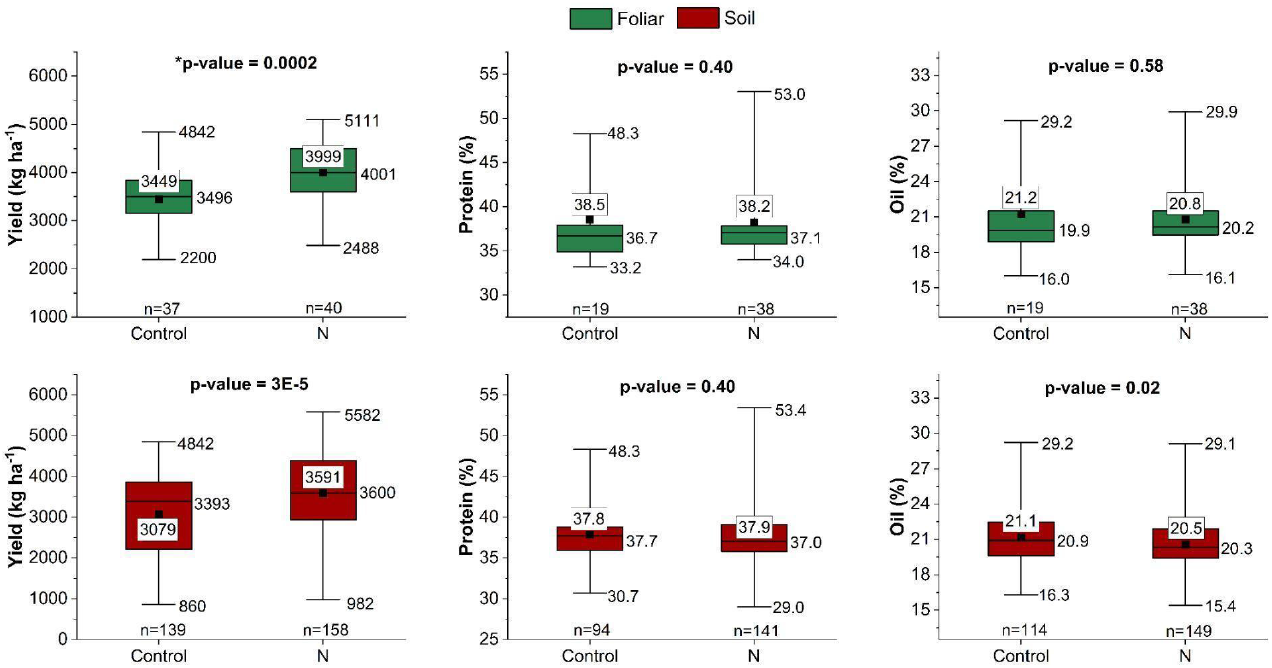
Comparison between N application and their control samples in foliar and soil application. *p-value from Mann-Whitney test.

Moreover, Figure 6 shows the correlation between different N doses by yield, protein, and oil content. Significant correlations (at 99% confidence level) were observed between applied N doses (both foliar and soil) with protein and oil contents. On the other hand, no correlation was observed between N doses and yield.

**Figure 6.**
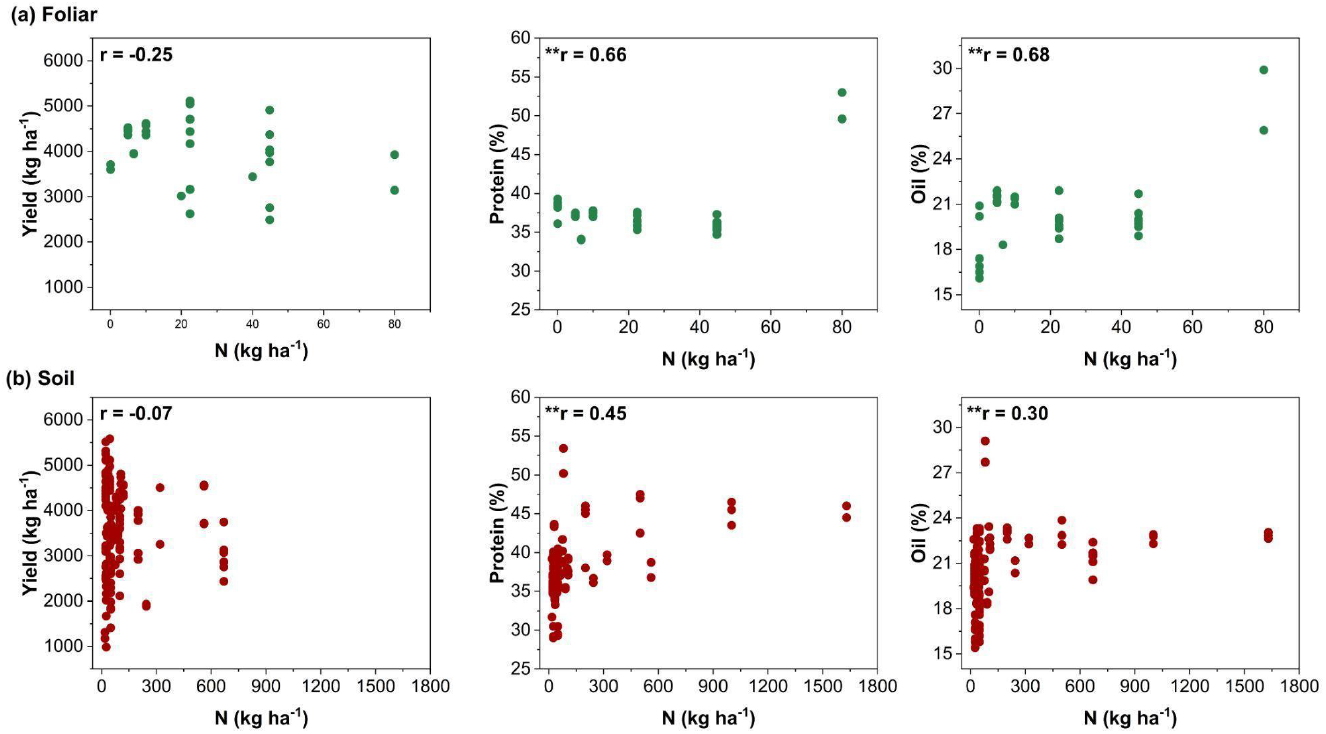
Correlation between N application dose, via (a) foliar and (b) soil fertilisation, and soybean yield, protein content, and oil content, with corresponding correlation coefficients. Statistically significant correlations at the 0.01 level are indicated by (**).

Due to the symbiotic association between legume roots and diazotrophic bacteria, primarily from *Bradyrhizobium* spp., soybean is commonly grown without nitrogen (N) fertilization (Bhangu, R., 2019). Nevertheless, numerous studies have investigated the effects of different N doses, timings, and application methods on soybean growth (Schmitt, M. A. et al., 2001; Mourtzinis, S. et al., 2018; Barker, D. W. et al. 2005). Despite the yield increase observed with soil-applied nitrogen doses (Figure 5), this practice is not widely recommended, as biological nitrogen fixation (BNF) delivers nitrogen directly to the plant more sustainably and efficiently, with reduced environmental impacts. Nitrogen use may be better justified through foliar application, as several studies suggest it leads to more consistent improvements in plant performance and greater nitrogen use-efficiency, increasing the grain yield by 13.7% according to the Figure 5.

## 3.0 CONCLUSIONS AND PERSPECTIVES

The results highlight significant variability, influenced by the complex and region-specific nature of agriculture, including soil type, climate, and topography. The lack of standardized experimental approaches and the limited number of studies further contribute to this variation. While field trials effectively validate technologies in real production scenarios, controlled environments may be more suitable for investigating fundamental physiological and metabolic processes.

A notable finding is the predominance of nitrogen fertilization methods, despite the role played by biological nitrogen fixation (BNF). Additionally, while nutrient application had no significant impact on protein and oil content, it led to a slight yield increase of 6.9%. The limited focus on foliar fertilization (22.9% of studies) underscores the need for further research on its potential to enhance soybean protein content. The trade-off between protein and oil content in soybean seeds warrants further investigation, as different nutritional strategies, such as applications of nitrogen or sulfur, can directly influence the biosynthesis of these biomolecules, potentially altering their balance within the seed.

This meta-data analysis revealed that the application of macro- or micronutrients, either via soil or foliar application, can directly influence plant yield. However, such increases in biomass do not necessarily translate into changes in protein and oil content, often resulting in higher grain production with similar concentrations of these biomolecules per unit area in field conditions.

In general, the results across the reviewed studies exhibited considerable variability and inconsistency, as well as the lack of clear methodological information, making it difficult to draw definitive conclusions regarding the effectiveness of nitrogen fertilization in soybean.

This study highlights the global relevance of fertilization practices in soybean production, demonstrating how different methods can influence plant responses and enhance yield across diverse environments. It also identifies key knowledge gaps, offering opportunities for future research aimed at deepening our understanding and improving agricultural practices.

## Supporting information

Dataset used for the literature review

## 4.0 ACKNOWLEDGMENTS

## 7.0 AUTHOR CONTRIBUTIONS

FSF: writing - review & editing; GPL: writing - review & editing; NGCS: data curation, writing - review & editing; JRB.: data curation, writing - review & editing; NLC: writing - review & editing; FRS: conceptualization, data curation, writing - review & editing; GSM: conceptualization, data curation, writing - review & editing; HWPC: conceptualization, data analysis, writing - review & editing.

## Electronic Supplementary Information

**Table S1:**
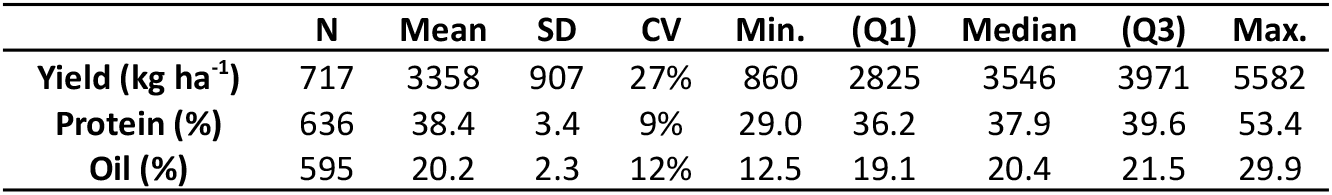
Descriptive statistics from the data of productivity, protein and oil.

**Table S2.**
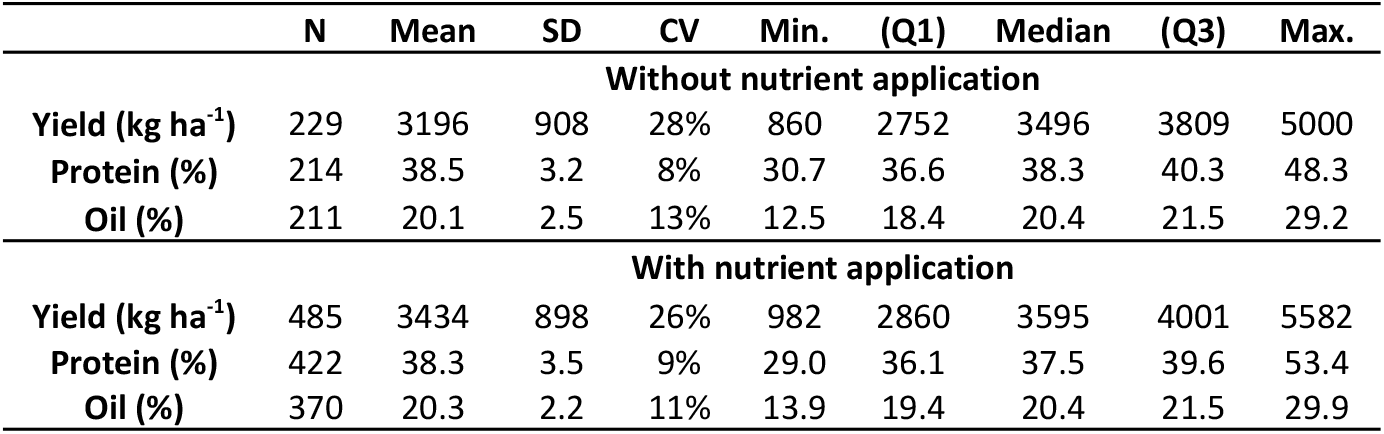
Descriptive statistics from the data of productivity, protein and oil content with and without nutrient application.

**Table S3:**
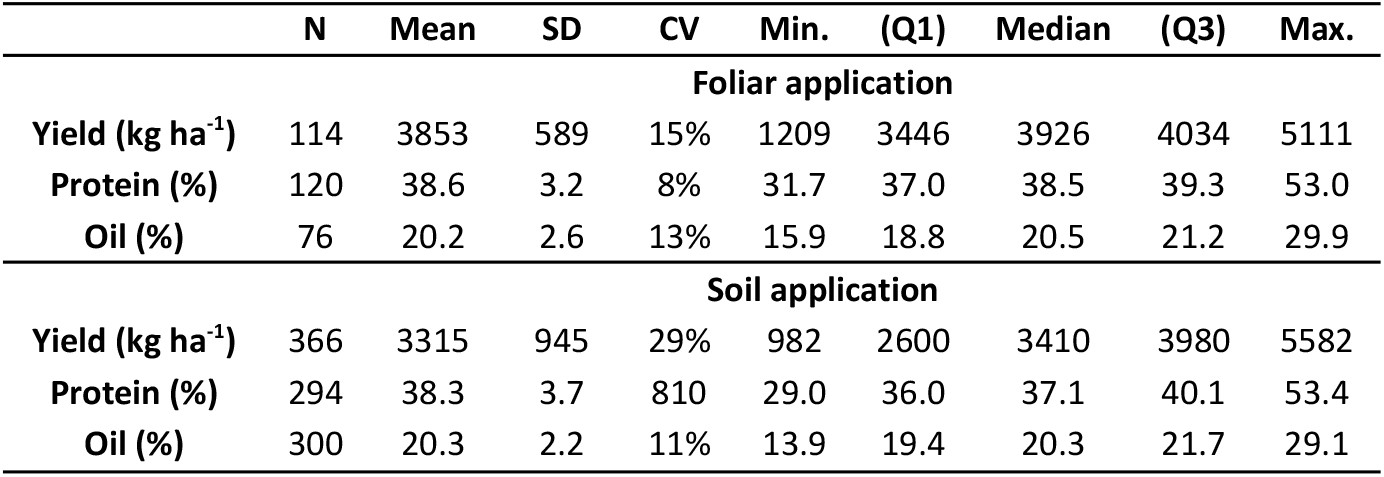
Descriptive statistics of productivity, protein and oil data reported with the application of nutrients via foliar and soil in the 48 studies analyzed.

**Table S4:**
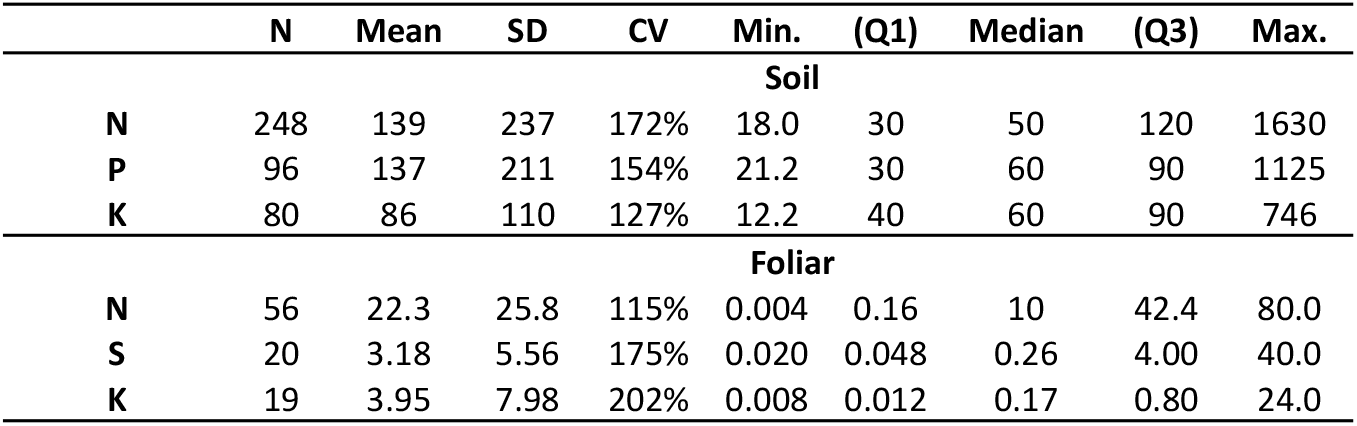
Doses of the main nutrients (Figure 4) applied via soil and foliar in the fertilization management strategy employed in the 48 studies analyzed.

**Table S5:**
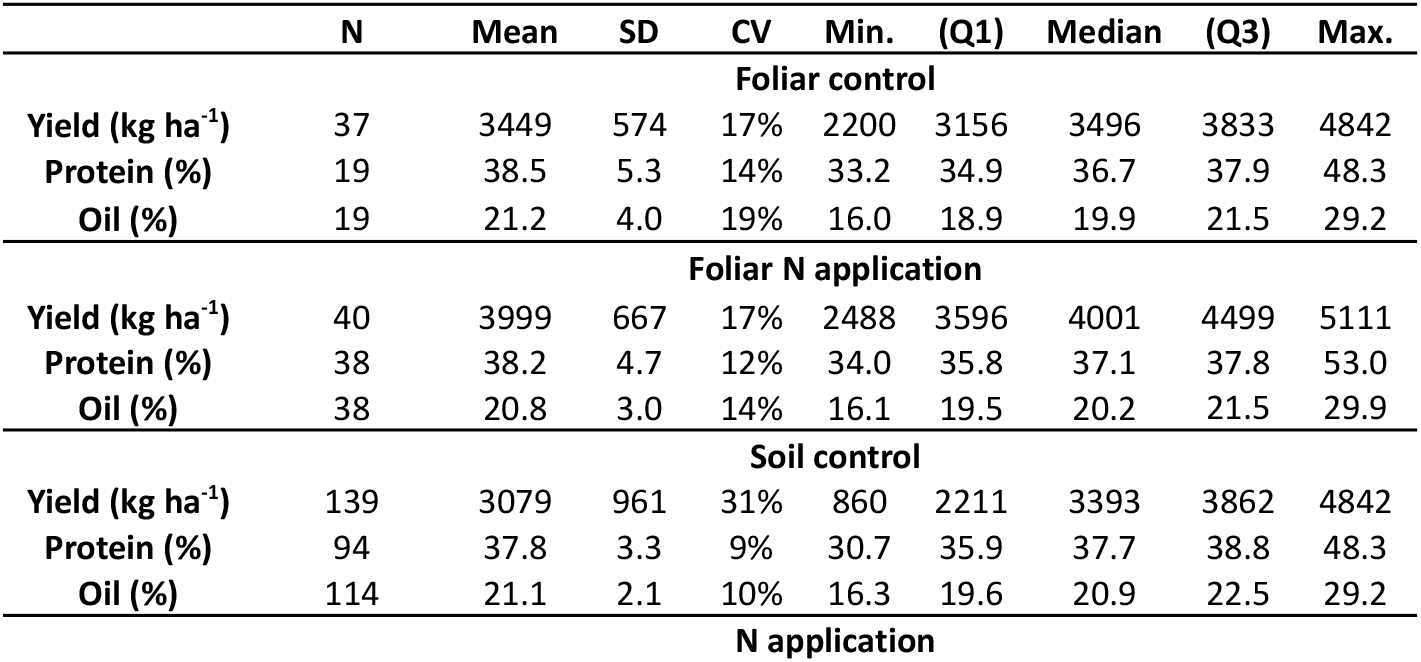
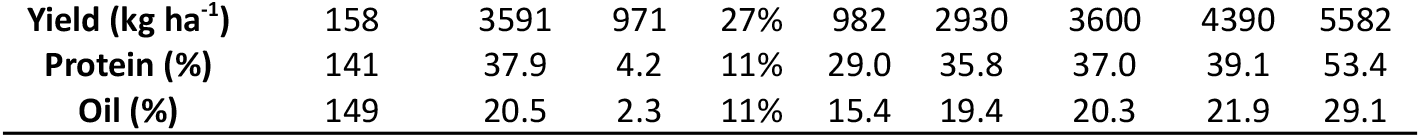
descriptive statistics of soybean yield, protein and oil content from the control samples and with N application through leaves and soil.

**Figure S1.**
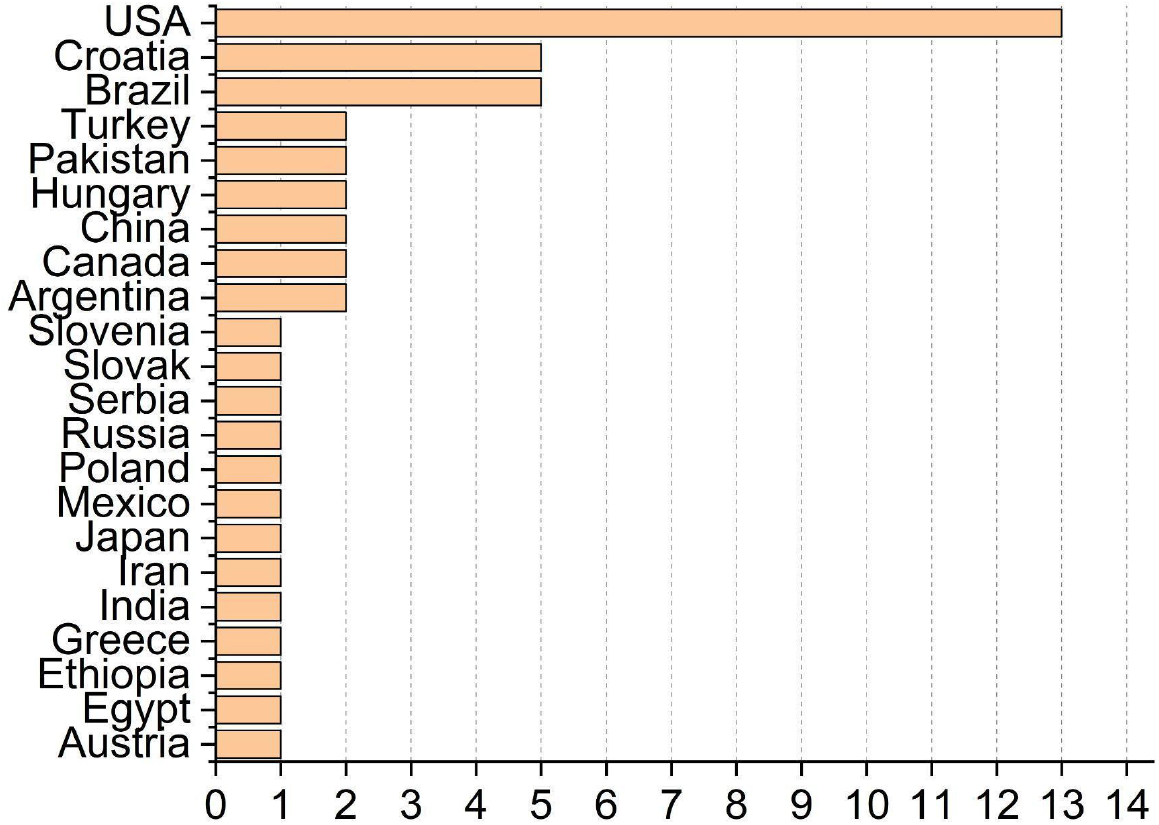
Number of studies per county in the 48 studies analyzed in the metadata analysis.

**Figure S2.**
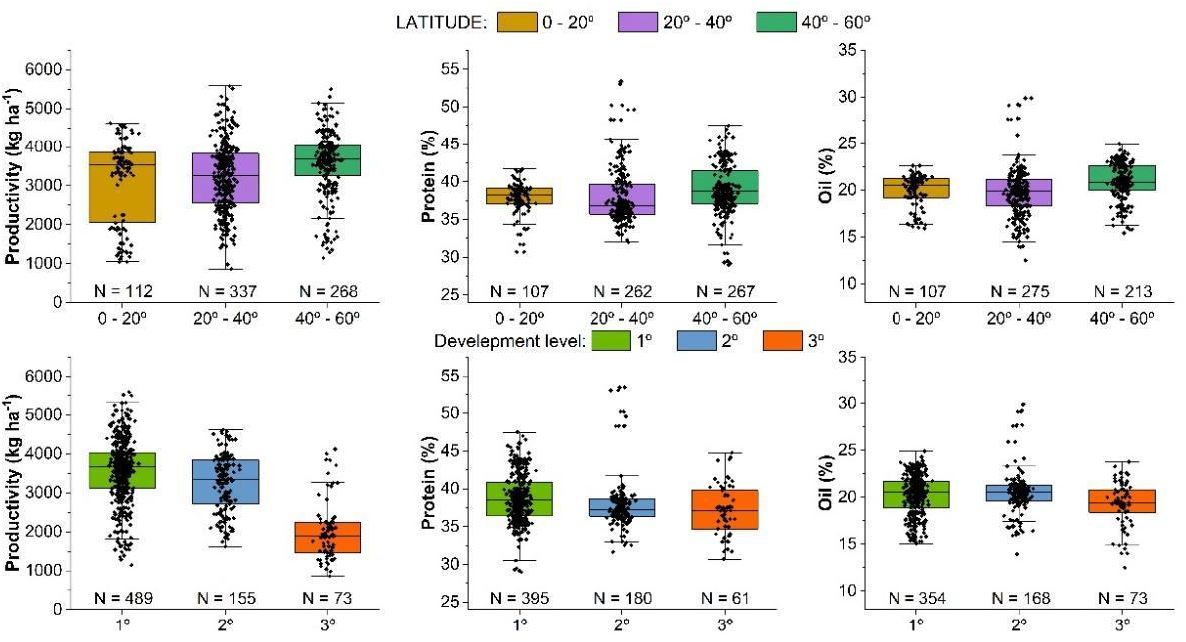
Productivity (Kg/ha), protein (%) and oil (%) content of soybean seeds according to latitude (Upper) and the level of development of countries (lower) where the experiments were conducted.

**Figure S3.**
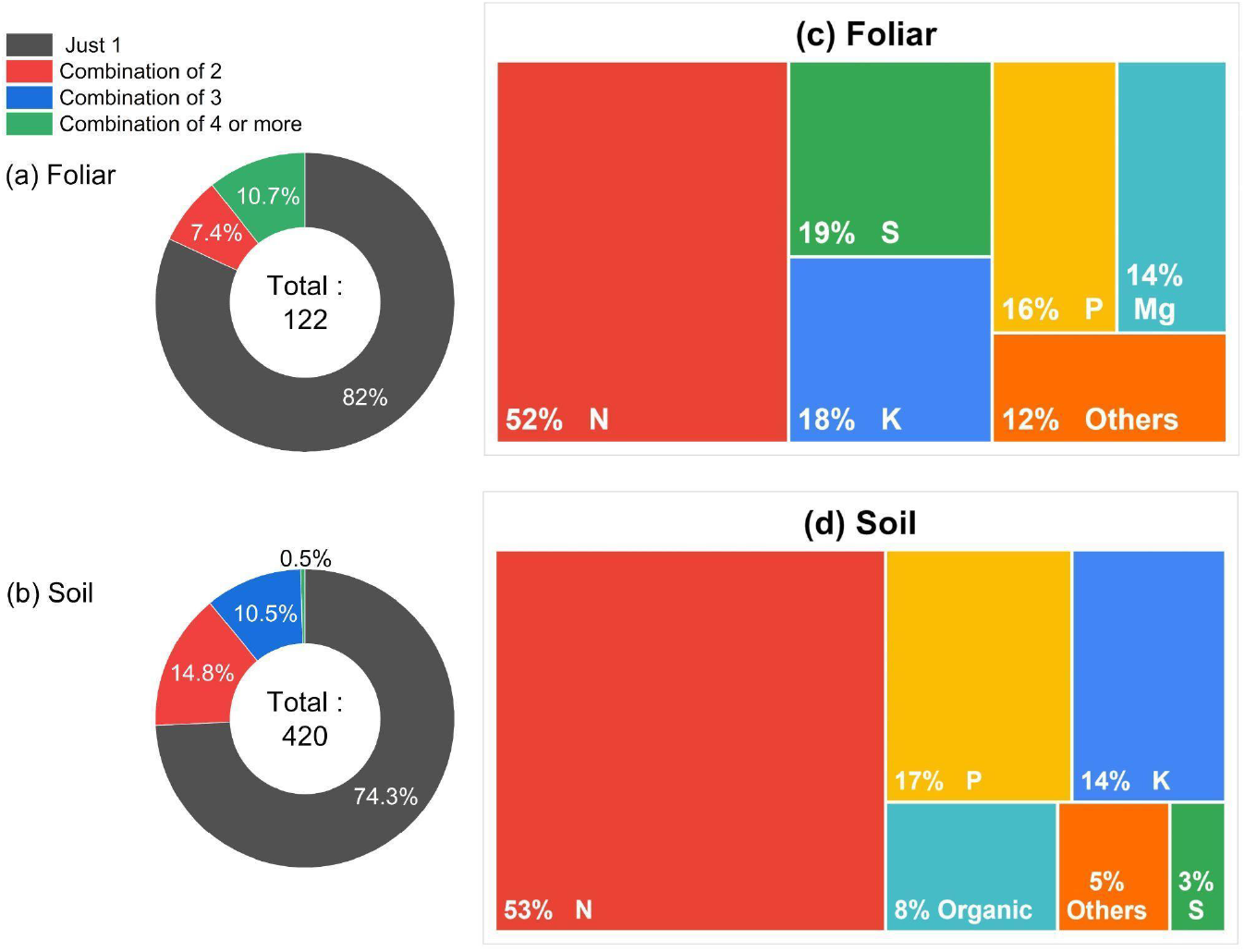
How many, which, and how *i.e*. soil or foliar, nutrients were employed in the fertilization management strategy of the 48 studies analyzed.

**Figure S4.**
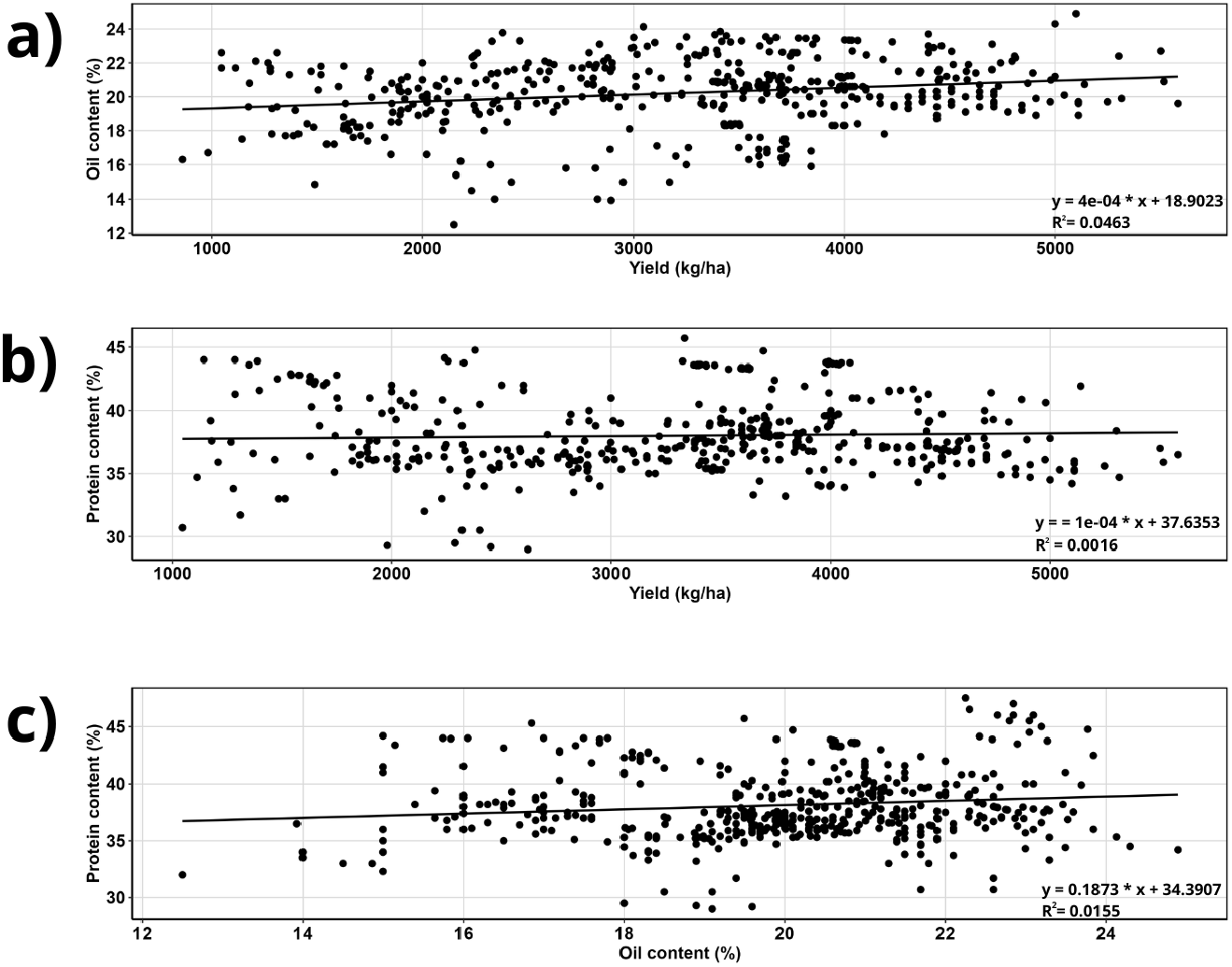
**Relationships between grain yield and oil content (a), grain yield and protein content (b), and between grain oil and protein concentrations (c) in soybean grains related to the 48 studies**.

### OUTLINE

#### 1. IMPORTANCE OF SOYBEAN AS PROTEIN SOURCE

*Importance of soybean proteins on animal and human nutrition*

*Temporal view on soybean cultivation*

##### 1.1 Challenges towards increasing protein content in soybean seeds

*Tradeoff between yield and protein content of soybean*

*Importance of mineral nutrition on soybean yield*

*Importance of mineral nutrition and protein synthesis in soybean*

#### 2. EFFECTS OF MINERAL NUTRITION ON SOYBEAN PROTEIN CONTENT: MEDATA OUTPUTS

*Experimental design and statistical analyses*

*Detailing fertilizer and application approach: Fig. 1 e Fig. 4*

*General Effects: Fig. 2, Fig. 3, Fig. 5*

*Effects of N: Fig. 7, Fig. 8*

## 3. CONCLUSIONS AND FUTURE DIRECTIONS

- limiting number of studies;
- importance of determining protein content;
- micronutrients;
- Last paragraph: there is plenty of room for boosting both the content and quality of soybean-based proteins by means of mineral nutrition.

